# Nucleolin internalizes *Bothrops asper* Lys49 phospholipase A_2_ forming cell surface amyloid-like assemblies

**DOI:** 10.1101/188383

**Authors:** Maria Lina Massimino, Morena Simonato, Barbara Spolaore, Cinzia Franchin, Giorgio Arrigoni, Oriano Marin, Laura Monturiol-Gross, Julián Fernández, Bruno Lomonte, Fiorella Tonello

**Affiliations:** Istituto di Neuroscienze, CNR, Via Ugo Bassi 58/B, 35131, Padova, Italy; Dipartimento di Scienze del Farmaco, Università di Padova, Via F. Marzolo, 5, 35131, Padova, Italy; Dipartimento di Scienze Biomediche, Università di Padova, Via Ugo Bassi 58/B, 35131, Padova, Italy; Centro di Proteomica, Università di Padova e Azienda Ospedaliera di Padova, Via G. Orus 2/B, 35129 Padova, Italy; Instituto Clodomiro Picado, Facultad de Microbiología, Universidad de Costa Rica, San José 11501, Costa Rica

## Abstract

Phospholipases A_2_ (PLA_2_s) are a major component of snake venoms. Some of them cause severe muscle necrosis through a still unknown mechanism. Phospholipid hydrolysis is a possible explanation of their toxic action, but catalytic and toxic properties of PLA_2_s are not directly connected. In addition, viperid venoms contain PLA_2_-like proteins, which are very toxic even if they lack catalytic activity due to a critical mutation in position 49. Nucleolin, a main component of the nucleolus, is a disordered protein involved in many protein assembly and phase separation phenomena. In some circumstances nucleolin is exposed on the cell surface from where it is involved in the internalization of many ligands.

In this work we demonstrate that *Bothrops asper* myotoxin II (Mt-II), a Lys49 PLA_2_-like toxin, interacts with, and is internalized in cells by nucleolin. The internalization process is functional to the toxicity of the protein, as both an antibody and an aptamer specific for nucleolin protect cells from intoxication. We identified central RRM and the C-terminal R/F-GG domain of nucleolin as the regions involved in the interaction with Mt-II. Finally we observed that Mt-II forms, on the cell surface, amyloid-like assemblies that colocalize with nucleolin and that can be involved in the activation of the internalization process. The presence, in the three dimensional structure of Mt-II and related PLA_2_ homologues, of four exposed loops enriched in prion-like amino acid sequences reinforces this hypothesis.

Phospholipases A_2_ | Lys49 myotoxins | nucleolin | amyloid-like | molecular assemblies

**SIGNIFICANCE:** The main finding of this work, the role of nucleolin as *Bothrops asper* Mt-II receptor, is a remarkable step forward in understanding the mechanism of action of cytotoxic PLA_2_s. It may suggest new strategies for anti-venom therapies and explain the anti-tumoral and anti-viral pharmacological action of snake PLA_2_s, since nucleolin is a receptor for many growth factors and virus.

The proposed internalization mechanism, via formation of molecular assemblies among Mt-II amyloid-like structures and other proteins, including nucleolin, can be of general validity. Cell surface molecular assemblies couldbepointsofselectionandconcentrationnotonlyofsnake,butalsoofmammaliansecretedPLA_2_s, proteins involved in different pathologies, and trigger the internalization pathway only when their molarity exceeds a threshold dose.

## INTRODUCTION

Snakebites affect every year more people than every other neglected tropical diseases, causing death but also permanent disability and disfigurement (*1*). Snake venoms consist of a mixture of toxins and enzymes that have evolved mainly to capture and digest the prey. Major clinical effects of envenoming in humans include coagulopathy, neurotoxicity, myotoxicity, and renal impairment, among others (*2*).

Phospholipases A_2_ (PLA_2_s) are major components of snake venoms, acting as hemostasis-impairing toxins, neurotoxins, or myotoxins. They have a high homology with mammalian secreted PLA_2_s, suggesting that they probably share cellular mechanisms and molecular interactors. This is of high relevance, in the light of the emerging involvement of mammalian secreted PLA_2_s in many human disorders (*3-5*). Various pharmacological applications have been proposed to exploit the diverse functionalities of snake PLA_2_s, including antiviral and antitumoral activities. However, the basic mechanisms of these activities are not known (*6-8*).

Most myotoxic PLA_2_s cause a local myonecrosis at the site of snakebite, but some of them act systemically, causing widespread muscle damage. Systemic myotoxins probably have high specificity for a muscle receptor, while locally-acting myotoxins, which induce myonecrosis only at relatively high doses, appear to interact with low-affinity receptors. Moreover, some local myotoxins also bind to and affect different types of cells, indicating that their receptor(s) is non-muscle-specific. Notwithstanding the many efforts made by several laboratories to identify myotoxic PLA_2_s receptors/acceptors in cell membranes, this search is still ongoing. Moreover, the internalization and possible interaction of these toxins with intracellular targets have not been explored (*7*).

A large subfamily of natural variants of snake PLA_2_s have no enzymatic activity, since they have a critical mutation at position 49: the aspartic acid is substituted by another amino acid (lysine in most cases), resulting in the impossibility to coordinate the calcium ion essential for catalysis. Despite the lack of catalytic activity, these PLA_2_ homologues show a high myotoxic activity and other toxic effects (*7, 9*).

*Bothrops asper* myotoxin II (Mt-II) is a Lys49 PLA_2_ homologue protein acting as a local myotoxin, but also affecting a wide variety of cell types *in vitro* (*10*), including macrophages (*11*). The currently held view is that Mt-II exerts its toxic activity by affecting the plasma membrane integrity, with consequent rapid influx of calcium ions that eventually triggers a series of degenerative events (*12*). However, this might be a simplistic view since the interaction of Mt-II with cells involves also intracellular signaling pathways: immediately after addition to cell cultures, Mt-II induces calcium release from intracellular stores, followed by opening of potassium and ATP channels, activation of purinergic receptors, and finally massive entry of calcium from extracellular medium (*11, 13, 14*).

In this work we conjugated Mt-II, purified from *Bothrops asper* venom, with a fluorophore to inquire its localization in target cells, and with biotin to use it as bait to isolate its protein interactors. We found that the toxin is rapidly internalized both in myotubes and in macrophages, and transported to the paranuclear and nuclear zone. Among the protein interactors we identified nucleolin (NCL), a multifunctional protein with a high percentage of disordered domains (*15*)and that is ubiquitously distributed in various eukaryotic cell compartments, such as the nucleolus, the nucleoplasm, the cytoplasm and the cell membrane (*16*). NCL has been reported to mediate the internalization of different types of molecules (*16, 17*). We verified the involvement of NCL in Mt-II internalization and toxic activity with pull-down experiments, cellular uptake, and cytotoxicity tests in presence of NCL competitors. Moreover, thanks to the different nature of these competitors, we were able to map the NCL domains involved in interaction with Mt-II.

Finally, we found that NCL co-localizes with Mt-II on the cell surface in structures that are sensitive to Congo Red staining. This observation, reinforced by the biochemical properties of NCL as disordered protein, and by the presence of prion-like sequences in the Mt-II structure, led us to hypothesize that Mt-II forms molecular assemblies on the cell surface that could trigger the internalization process and are probably also involved in cell membrane permeabilization.

## RESULTS

### Mt-II is internalized in myotubes and macrophages

Purified Mt-II was conjugated, by reaction with transglutaminase (*18*), to TAMRA and DNS fluorophore containing peptides to observe the localization of the toxin in target cells, myotubes and macrophages. The major reaction product was isolated by RP-HPLC and characterized by ESI-mass spectrometry and by cytotoxicity test. The determined molecular masses correspond to that of the mono-conjugated products and the toxic activity is, for more than eighty percent, conserved (see supplementary Fig. S1).

Mt-II resulted to be rapidly internalized (within minutes) both in myotubes and in macrophages in an asynchronous way: some cells are intoxicated before others, and cell death is preceded by the internalization of the toxin (Fig. 1 and Video S1, S2 and S3). The internalized toxin is transported toward the cell nucleus and in some cases co-localizes with it or with nuclear membrane that appears prominent in dying cells, while no important colocalization with mitochondria was observed (Fig. 1A). Time lapse experiments of macrophages and C2C12 myotubes treated with Mt-II DNS (Video S1 and S3) evidence that the toxin induces membrane blebs typical of macropinocytosis (*11, 19*) and that higher doses of Mt-II cause a rapid detachment of the cells from the substrate (Video S2).

**Fig. 1.**
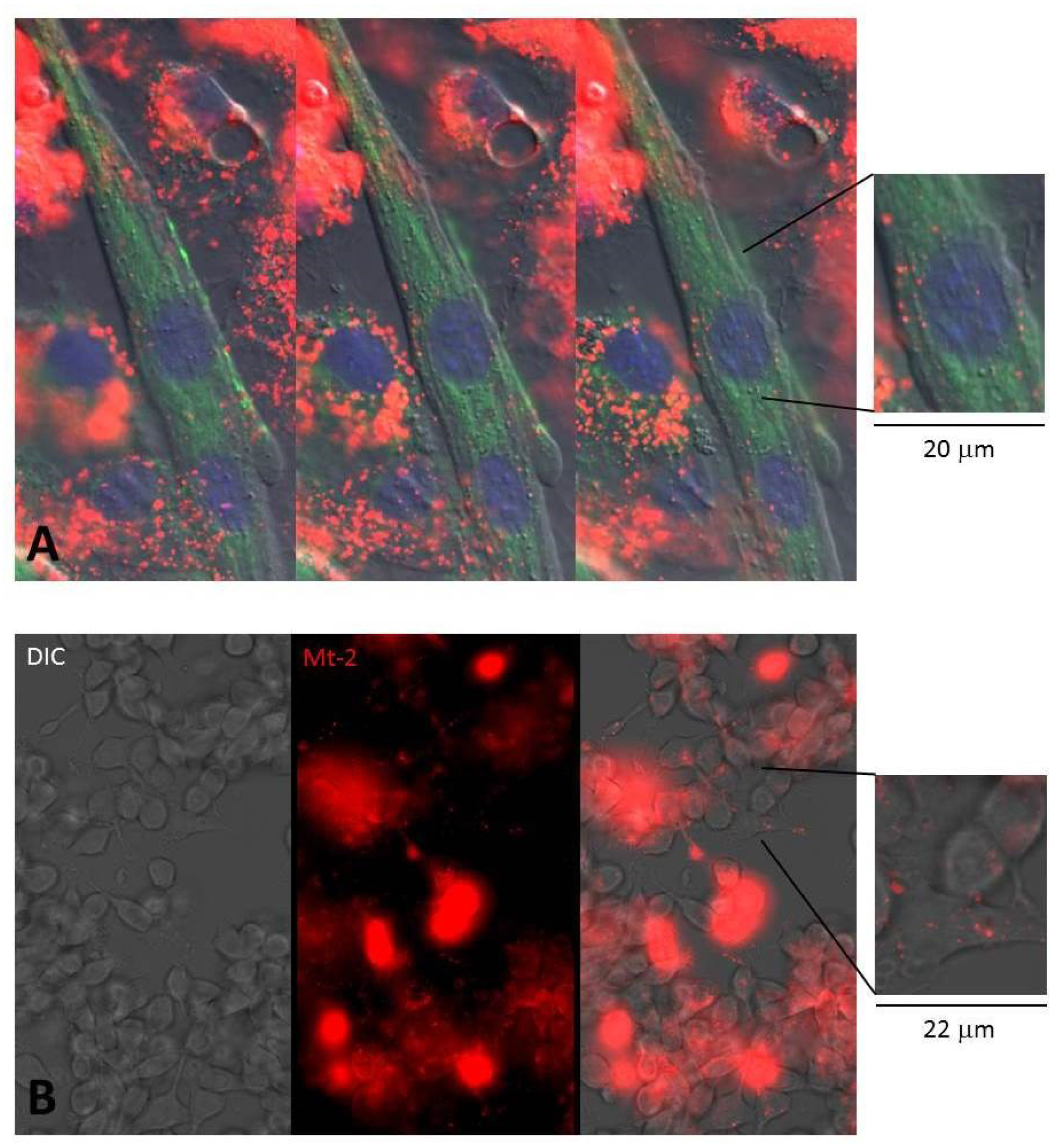
Internalization of Mt-II in myotubes and macrophages. **A**. Mouse primary myotubes intoxicated for 30’, 37°C, with 1 μM Mt-II-TAMRA (red). Mitochondria were stained with MitoTracker Green (1:2000) and nuclei with Hoechst 33342 (1:1000). Three observation planes (4 μm total thickness) were reported. **B**. RAW264.7 cells were treated for 30’, 37°C with a 1 μM Mt-2-TAMRA (red) washed and observed. Images were acquired with wide-field epifluorescence and transmitted light microscope (DIC Nomarski). Both cell types fill with toxin before dying, in an asynchronous way as if some cells are more sensitive than others. In the pictures intact cells containing few Mt-II containing vesicles (see inserts), are surrounded by damaged cells filled with the toxin.

### Isolation and identification of Mt-II protein interactors from macrophage cell extracts

Mt-II was conjugated to a biotin containing peptide by reaction with transglutaminase. The reaction and the protein purification and characterization were executed as reported in the previous paragraph. Biotinylated Mt-II (Mt-II-B), was combined to streptavidin magnetic beads and utilized as bait to pull-down interacting proteins from a RAW264.7 cell extract. The isolated proteins were eluted with a 5% sodium deoxycholate (SDC) solution, that does not detach the myotoxin from the resin, then digested with trypsin and identified by LC-MS/MS analysis. As controls, the same procedure was performed using beads without Mt-II. The experiment was repeated twice under the same conditions, each time leading to the identification of about one hundred proteins, each of them with a false discovery rate (FDR) less or equal to 0.01, and with at least four sequenced unique peptides (Table S1 and S2). To identify proteins closer to the bait, the experiment was then repeated a third time by adding a crosslinking step on streptavidin beads, after protein isolation. The crosslinker (DTSSP) contains amine-reactive NHS-ester ends around a 12 Å, 8-atom spacer arm, and a central disulfide bond that can be cleaved with reducing agents. After the crosslinking reaction the non-covalently bound proteins were removed with the 5% SDC solution and finally crosslinked proteins were detached with 50 mM DTT dissolved in the 5% SDC solution. Also in this case about a hundred proteins were identified but with a lower abundance: only 21 proteins with a sample/control intensity ratio higher than 3 (Table S3), in comparison to 111 and 84 of the first and second experiment. Fifteen proteins were found in common in all three experiments (Fig. 2). NCL was identified in all pull-downs among the proteins with the largest number of unique peptides and with an estimated abundance at least 45 times higher with respect to the control sample. Since NCL is a nucleolar protein known to be present also on the cell surface as well as a receptor or co-receptor of different factors, we decided to enquire the Mt-II/NCL interaction and the role of NCL in Mt-II internalization.

**Fig. 2.**
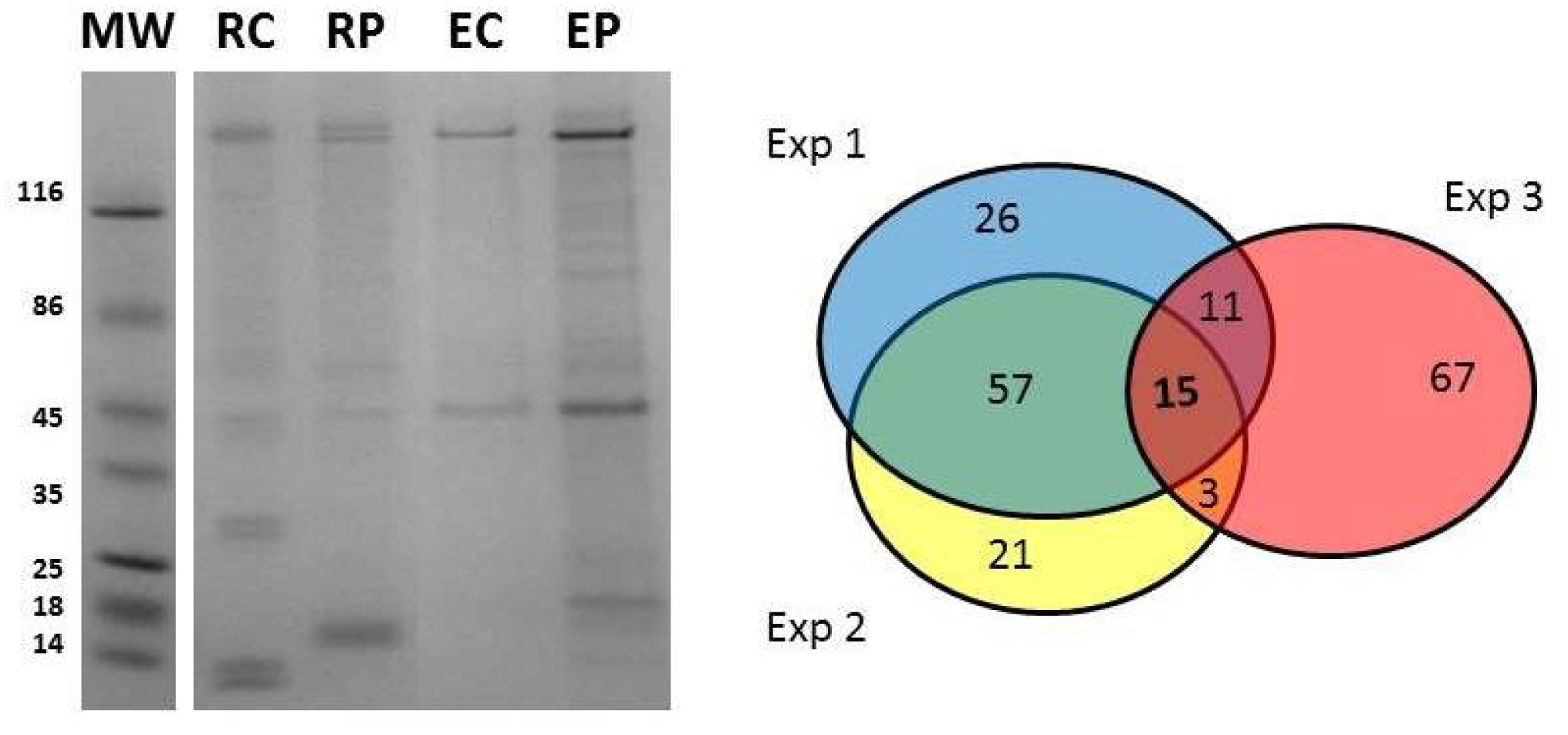
Pull down and mass spectrometry analysis of Mt-II protein interactors. **(Left)** SDS-PAGE analysis of pull down experiment with biotinylated Mt-II (Mt-II-B) combined with streptavidin magnetic beads (SB) as bait and an extract of RAW264.7 as prey. As control the pull down was also executed with the resin without Mt-II-B. Pulled down proteins were eluted with a solution containing 5 % SDC in 50 mM NH_4_HCO_3_. **RC**: SB after elution; **RP**: SB-Mt-II-B after elution of pulled-down proteins; **EC**: sample eluted from SB; **EP**: sample eluted from SB-Mt-II-B. Samples **EC** and **EP** were subjected to in-gel digestion followed by LC–MS/MS and semi-quantitative data analysis (Table S1-S4). The experiment was repeated two times with the same conditions, and a third with the addition of a crosslinking step to isolate proteins closer to the bait. (**Right**) Venn diagram of the number of proteins identified in the three pull down experiments. Among the fifteen proteins present in all three experiments NCL was identified with at least 13 unique peptides and an intensity ratio, sample over control, higher than 45.

### Mt-II pulls down nucleolin from RAW264.7, C2C12 and *ex-vivo* muscle membrane preparations

NCL is one of the most abundant non-ribosomal proteins of the cell. It is mostly localized in nucleolus (90%) and nucleus (about 5%), but it is present also in cytosol and cell membrane in variable concentrations, depending on cell status (*16, 20*). To confirm the interaction between Mt-II and NCL in cell types other than macrophages, and to verify that Mt-II interacts with NCL present in cell membrane, we prepared an extract of membranes from RAW264.7, C2C12 myotubes and from *ex-vivo* muscle. The subcellular fractionation was performed with a commercial kit and the fractions were characterized by western blot to verify the efficiency of the protocol (Fig. S2). Membrane fractions were incubated with Mt-II-B combined with streptavidin magnetic beads, eluted by 5% SDC solution and analyzed by western blot. As observed in Fig. 3 A, Mt-II pulls down NCL from membrane extracts obtained from all three preparations.

**Fig. 3.**
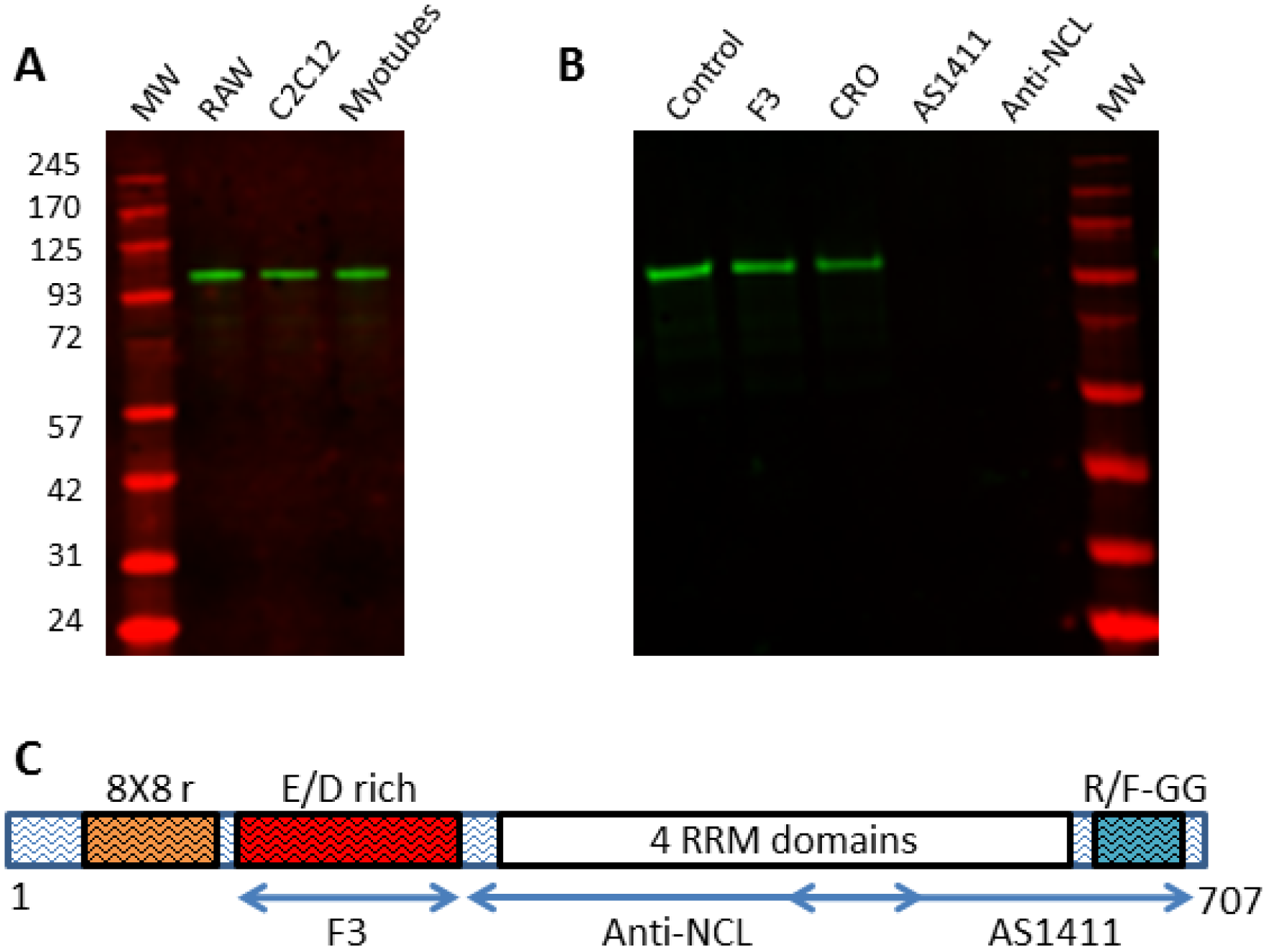
Mt-II pulls down NCL from membrane protein extracts, by interacting with NCL central and C-terminal regions. Western blot analysis, marked with anti-NCL, of proteins pulled down by Mt-II-B/streptavidin magnetic beads from (**A**) RAW264.7, C2C12 and mouse primary myotubes membrane protein extracts; (**B**) RAW264.7 membrane protein extracts alone (control) or in presence of NCL binding molecules: a peptide (F3, 5 μM), an aptamer (AS1411, 5 μM), and an antibody (anti-NCL, 20 μl). CRO is an aptamer used as control (5 μM). (**C**) Representation of the primary structure of mouse NCL and its domains. Regions recognized by the different antagonists used in this work were highlighted with double-headed arrows.

NCL is a protein of 710 (human) or 707 (mouse) amino acids, respectively, composed of three domains: an N-terminal disordered domain rich in negatively charged amino acids, a central domain containing four RNA Recognition Motifs (RRMs), and a C-terminal disordered domain containing R/F-GG repeats (Fig. 3 C). Many substances were found to interact with NCL present on the cell surface, in particular on surface of cancer cells where NCL is present at higher concentration, and to trigger an internalization process (*17, 20*). AS1411 is an anticancer aptamer that binds to the central and C-terminal regions of NCL (*21*), and F3 is a tumor-homing peptide that binds to the N-terminal negatively charged domain of NCL (*22*). We tested the ability of AS1411, F3 and of an anti-NCL central domain polyclonal antibody to compete for the pull-down of NCL by Mt-II-B. Interestingly, the antibody and the aptamer inhibit the pull-down of NCL while F3 does not compete (Fig. 3 B), indicating that the central and C-terminal regions of NCL are involved in the interaction with Mt-II.

### AS1411 and anti-nucleolin antibody inhibit Mt-II cell internalization and toxic action

To understand if the interaction between NCL and Mt-II has a role in the internalization and toxic activity, we measured the quantity of Mt-II internalized and the cytotoxicity of the protein in the presence of AS1411, a control aptamer (CRO), and an anti-NCL antibody. We performed an internalization assay intoxicating target cells with Mt-II-TAMRA, washing extensively to remove unbound protein, and measuring the fluorescence after cell lysis. For the cytotoxicity assay, cells were incubated with unlabeled Mt-II and the percentage of cell death was measured with a colorimetric assay for assessing cell metabolic activity, in the case or RAW264.7 cells, or by measuring the release of LDH, a plasma membrane damage index, in the case of the myotubes. The results of the two assays are reported in Fig. 4: the two NCL competitors significantly inhibited, though not completely, both the internalization and the toxicity of Mt-II.

**Fig. 4.**
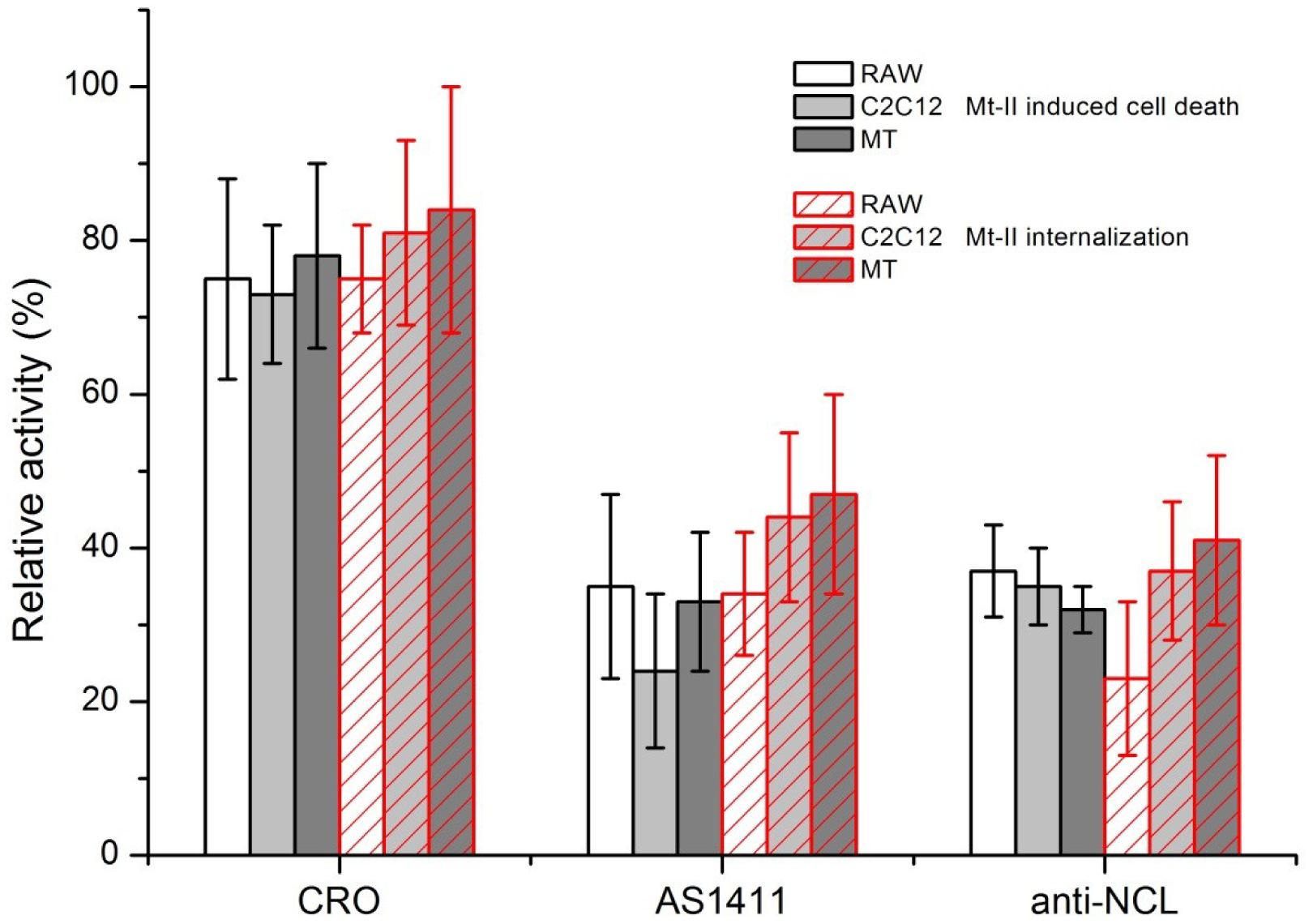
Inhibitory activity of NCL antagonists in cells intoxicated with Mt-II. RAW264.7, C2C12 and mouse primary myotube (MT) cells were treated with 1.5 μM Mt-II for 30’ at 37°C or with 1 μM Mt-II TAMRA for 10’, 37°C and checked for their vitality or for the internalization of the fluorescent toxin. The experiments were repeated in presence of two of the nucleolin antagonists, described in Fig. 3, same quantities, and the percentage of residual activity was calculated respect to control (100%). The plot represents the mean of three independent experiments, bars represent standard deviations.

### Mt-II co-localizes with nucleolin in cell surface assemblies sensitive to Congo Red staining

The interaction between NCL and Mt-II was verified also by co-localization experiments. In developing the experimental protocol, we observed that the amount of NCL present in the cell surface of non-stimulated cells is very low, but it increases in live cells treated with the toxin or with an anti-NCL antibody, at RT or 37°C. In fact NCL is released to the surface through an unconventional secretion that does not pass through the classic ER-Golgi pathway (*23*). Since we aimed at observing the interaction between NCL and Mt-II on the cell surface, we decided to add the toxin at low temperature (4°C) to prevent its internalization and consequent cell death. Thus, we treated cells with the anti-NCL antibody at RT to stimulate the secretion of NCL and successively with Mt-II-TAMRA at 4°C. Finally, we fixed the cells, treated them with the secondary antibody and we acquired images by confocal microscopy. The obtained images (Fig. 5 and Fig. S3) show that NCL and Mt-II colocalize in long stretches, fairly thick, on the cell surface. This kind of staining, non-dotted but with bigger areas, is typical of proteins involved in phase transition phenomena that give rise to membrane-less organelles (*15, 24*). Membrane-less organelles are molecular assemblies that form through multivalent weak interactions and that mediate diverse biological processes. The nucleolus is an example of this kind of organelles and NCL is one of its main components (*25*). Phase transition phenomena can happen also on the cell membrane, for example to trigger signaling events (*26*). As secreted PLA_2_s were reported to form amyloid-like fibrils when they come in contact with phospholipid bilayers (*27*), and prion-like interactions are involved in phase transition phenomena, we decided to assess if Mt-II assemblies on cell membrane are sensitive to Congo Red, an amyloid specific dye. We observed polymer-like green birefringence on polarized light, on the surface of cells intoxicated with Mt-II (Fig. 6), indicative of the presence of amyloid-like fibrils, and red coloured zone that co-localize with NCL. Assemblies formed by Mt-II are not SDS-resistant: SDS-PAGE analysis of extracts from cells incubated with Mt-II-B and a membrane impermeable crosslinking agent shows that Mt-II forms high molecular weight complexes only in crosslinked samples (Fig. S4). Since proteins that form amyloid-like fibrils are characterized by the presence of prion-like domains (*15, 24*), we inspected the primary structure of Mt-II for the presence of prion-promoting amino acids. We found four traits rich in tyrosine, serine and glycine with some asparagine or glutamine, amino acids enriched in human prion-like domains (*28*), with in addition the presence of proline and charged amino acids that are relevant to modulate the formation of prion aggregates (*28, 29*). Interestingly, these traits are located in exposed loops of Mt-II (Fig. 7 B) and homologous proteins, sites that can undergo conformational changes in an otherwise very stable tertiary structure due to presence of seven disulphide bonds.

**Fig. 5.**
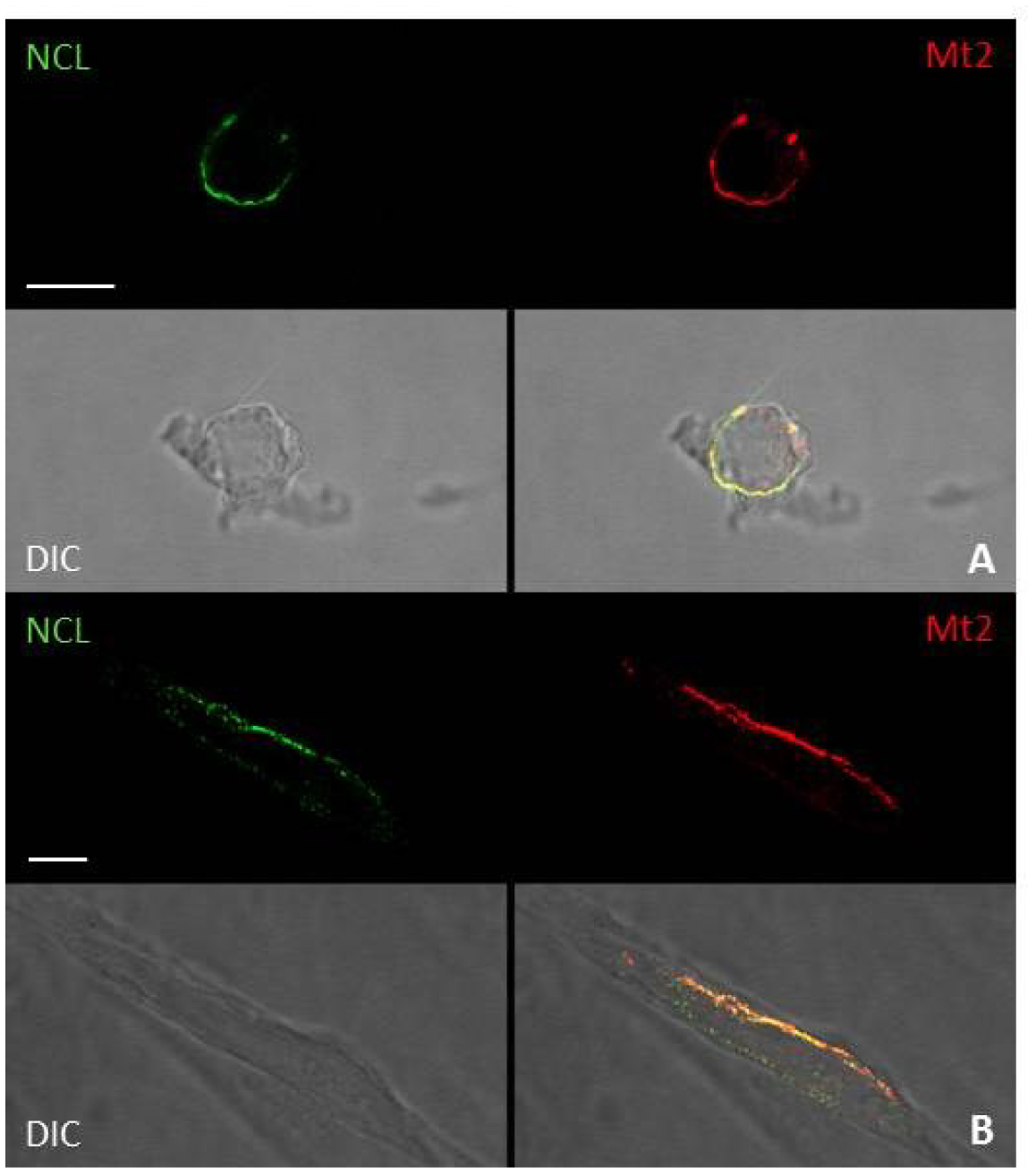
Colocalization of NCL and Mt-II in cell membrane of RAW264.7 and mouse primary myotubes. (see also Fig. S3 and Video S4). Scale bars correspond to 10 μm. RAW264.7 cells (**A**) and mouse primary myotubes (**B**) were incubated with anti-NCL (rabbit) (45’, RT), then treated with 1 μM Mt-II-TAMRA (15’, 4°C). After extensive washings in PBS, cells were fixed, treated with an anti-rabbit Ig secondary antibody and visualized by confocal microscopy.

**Fig. 6.**
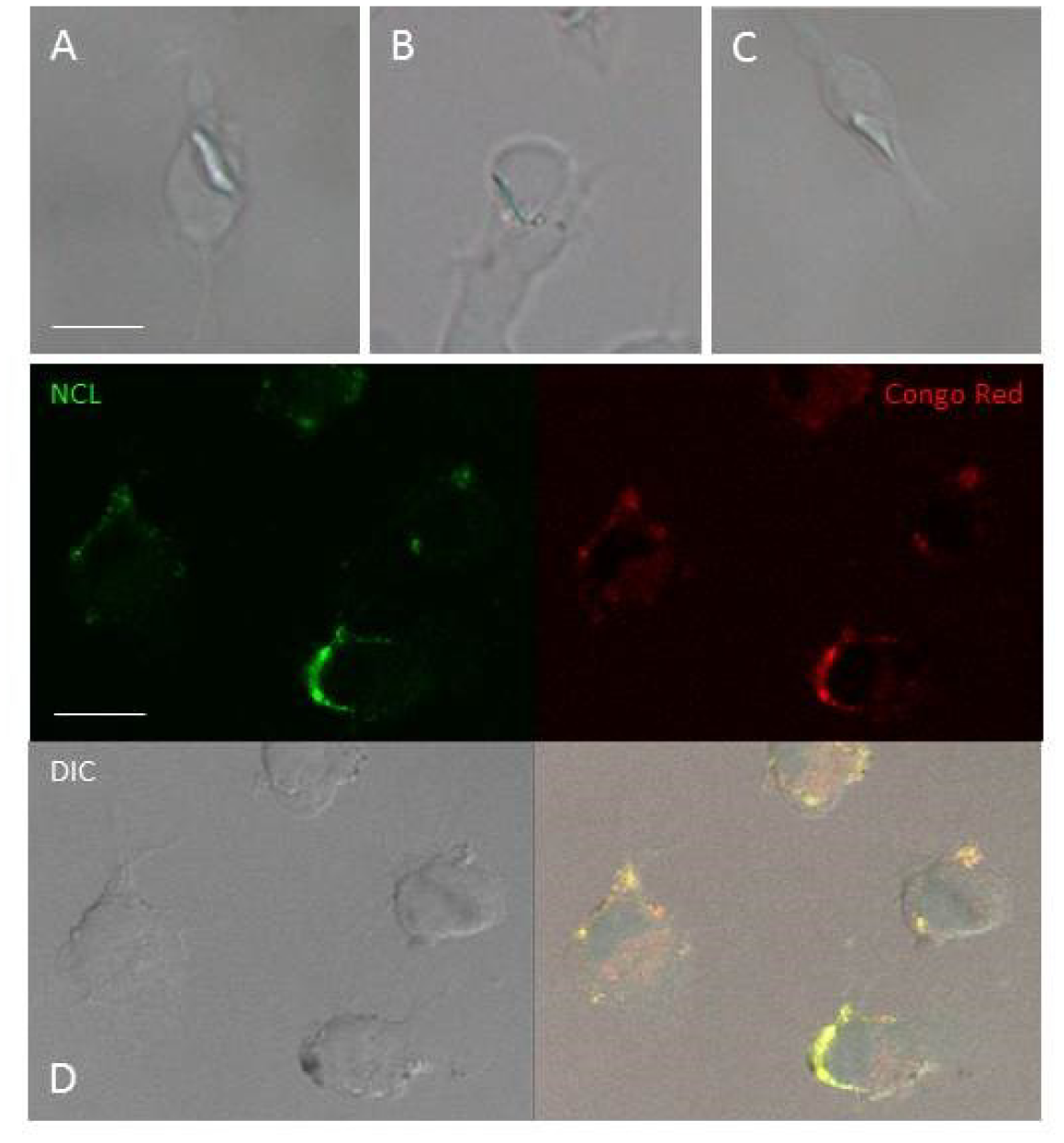
Congo red staining of Mt-II assemblies in cell membrane. RAW264.7 cells were treated with Mt-II (20’, 4°C), colored with Congo Red (10’, 4°C) fixed and observed in polarized light (**A-C**) or treated with rabbit polyclonal anti-NCL (45’, RT), with Mt-II (20’, 4°C), colored with Congo Red, fixed, treated with an anti-rabbit secondary antibody (green) and observed with a confocal microscope (**D**). Mt-II forms cell surface assemblies that, after Congo Red staining, show a green birefringence in polarized light and a red color at visible light (λ_exc_= 543 nm) that colocalize with the staining of NCL.

**Fig. 7.**
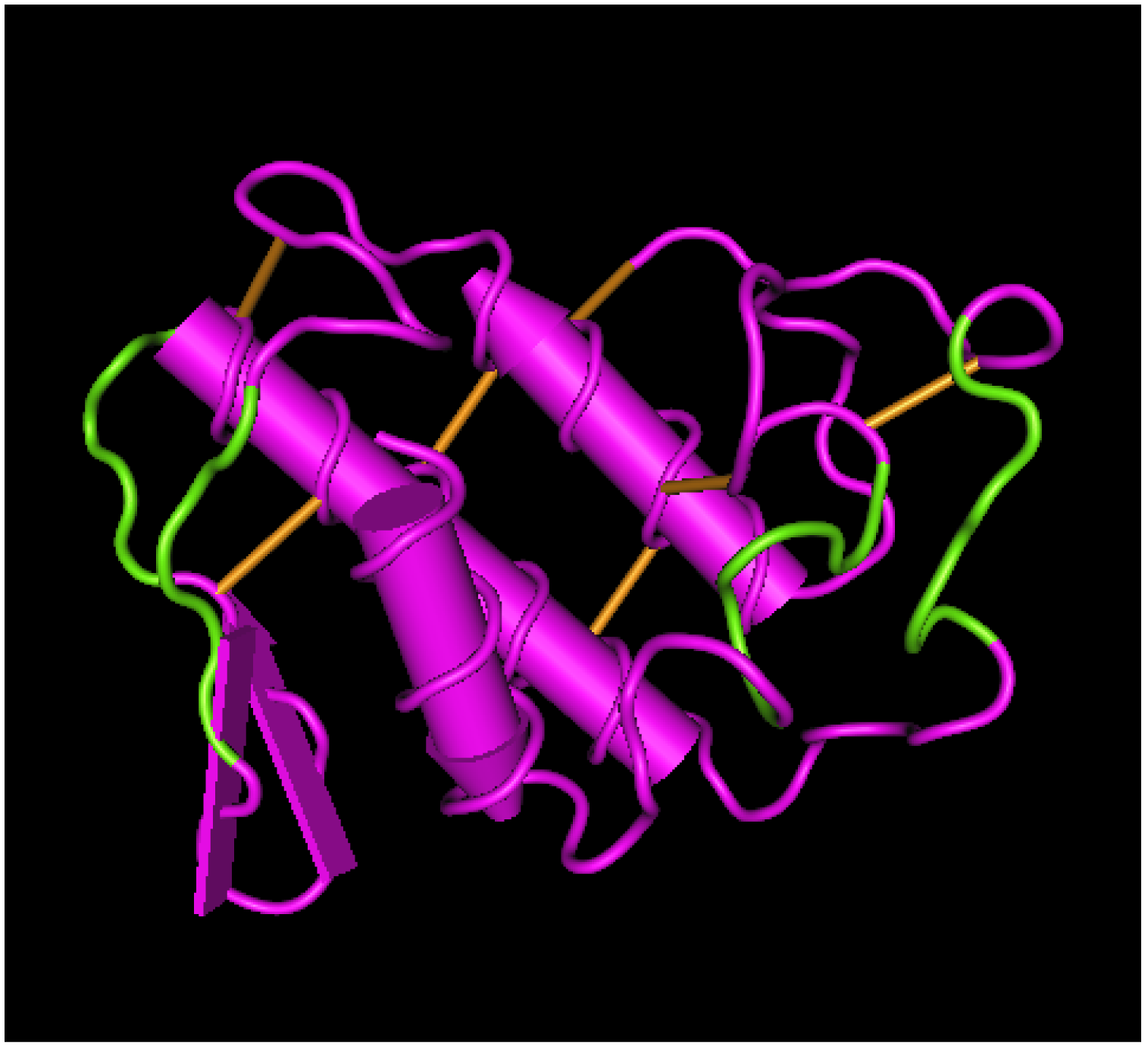
Presence of prion-like sequences on exposed loops of Mt-II and PLA2G2a. (**Up**) Alignment of primary sequence of *B. asper* K49 myotoxin (Mt-II) and D49 myotoxin (Mt-I), human and mouse PLA2G2a. Secondary structure elements were underlined and traits rich in prion-like amino acids were evidenced in green. (**Down**) Mt-II 3D model (PDB: 1CLP) where the traits rich in prion-like amino acids (see main text for definition) were colored in green. The picture was obtained with the Cn3D macromolecular structure viewing program.

## DISCUSSION

The first result of this work is the demonstration that *B. asper* Mt-II is internalized in myotubes and macrophages and that it localizes in perinuclear and nuclear zone (Fig. 1). Other snake PLA_2_s have been reported to be internalized, but only the neurotoxic PLA_2_s, and exclusively in neuronal cells. Notably, notexin, beta-bungarotoxin and taipoxin have been shown to localize in mitochondria of spinal cord motor neurons and cerebellar granule neurons (*30*), while ammodytoxin was found into the cytosol, synaptic vesicles, and mitochondria of motoneurons (*31, 32*). Snake cytotoxins, non-enzymatic three-fingered fold proteins, have been reported to be internalized in cells, and to induce cell death mainly by interaction with lysosomes (*33*). Importantly, PLA2g2a, the human secreted PLA_2_ with higher homology to Mt-II, was also reported to be internalized in monocytes and finally transported into the nucleus (*34*). Accordingly, the PLA2g2a subcellular localization reported in <u>Human protein atlas</u> (*35*) is nucleus and nucleolus.

The second result of this work is the isolation of putative cellular proteins interacting with *B. asper* Mt-II. NCL, one of the proteins identified in all pull-down/mass spectrometry experiments, is known to be present also on the plasma membrane and acts as a receptor or co-receptor for several factors (*16, 17*). The interaction of Mt-II with this protein was confirmed by pull-down/western blot experiments from cell membrane extracts and by the competition exerted by an anti-NCL antibody and an aptamer specific for NCL. The inhibition of cytotoxicity and internalization by these competitors was partial, but this result was expected considering that AS1411 and anti-NCL antibody, after internalization, induce the secretion of further NCL in a continuous process (*23, 36*). The Mt-II/NCL interaction is remarkable for several reasons. First, NCL is the first Mt-II interacting protein connected to the toxic mechanism. An interaction of Mt-II with the KDR/VEGF receptor 2 has been previously reported (*37, 38*) but no evidence for a functional role of this interaction was found by using a blocking monoclonal antibody to the receptor or the inhibitor tyrphostin (*9*). Second, the interaction of Mt-II with NCL may explain pharmacological properties of snake myotoxins. The preferential activity of these toxins against cancer cells (*6, 39, 40*) can be correlated to the fact that NCL is more abundant on the surface of these cells (*16, 41, 42*). Moreover, some snake PLA_2_s have been reported to have antiviral activity (*43, 44*) and accordingly NCL is involved in internalization of many viruses (*16, 17*). Third, several phenomena observed in cells following Mt-II intoxication can be explained by the interaction My-II/NCL: calcium entry is triggered also by NCL (*45*); the internalization of AS1411 takes place through macropinocytosis (*46*); NCL is involved also in cell adhesion and migration (*42*) and this could explain the rapid detachment of myotubes intoxicated with high doses of Mt-II (Video S2).

In the last part of this work we verified the colocalization of NCL and Mt-II in large areas of the cell surface and we observed that Mt-II forms amyloid-like structures in contact with cells. This suggests that the NCL-Mt-II interaction is not one-to-one but a multi-molecular assembly where other components are likely to be present. Nucleophosmin for example, a known partner of NCL, was identified among proteins co-precipitated by Mt-II (Table S4). However transmembrane proteins also can be involved, and they may not have been detected in our pull-down because they are more difficult to isolate and identify by mass spectrometry. In this regard it is worth remembering that both NCL and PLA2g2a interact with integrins and EGFR (*47-50*). Since NCL is not a membrane protein, it will have to interact with transmembrane proteins to communicate with the cell interior and trigger the internalization process.

NCL sequence (Fig. 3C), rich in disordered regions and low complexity domains, is typical of proteins participating in multimolecular interactions and phase transition phenomena, that is, transitions from dispersed protein solutions to liquid-like phase-separated compartments or to solid protein aggregates (*15, 24*). Mt-II, like other secreted PLA_2_s, has a well-defined and compact 3D structure stabilized by seven disulphide bonds; as a consequence, one would not expect it to be inclined to form multimolecular interactions. However, the fact that Mt-II forms amyloid-like structures, when in contact with cell surface, changes the view. Several proteins involved in phase transition phenomena possess prion-like domains, and prion structures contribute to the protein assembly. Moreover, many disordered proteins are thought to recognize and interact with amyloid structures, and NCL could be one of them. In fact, NCL is also a receptor of amyloid beta peptide 1-42 (*51*) and the R/FGG domain, that we found to be involved in interaction with Mt-II, is similar to the glycine rich domains of the chaperones involved in yeast prion propagation (*52*).

The formation of amyloid structures and multimolecular assemblies explains many characteristics of Mt-II and similar proteins. One is the need for a relatively high concentration of protein for toxic activity: locally-acting PLA_2_ myotoxins act at a concentration of micromolar order, while neurotoxic PLA_2_s and systemic myotoxins at a concentration even 1000-fold lower (*12*). Another is the tendency of Mt-II and other PLA_2_s to be unstable *in vitro*: when dissolved, we keep the protein in 50% glycerol otherwise it slowly forms aggregates, even at low temperatures. Finally, we should consider the antimicrobial capacity of Mt-II, and other snake PLA_2_s (*8, 53*) and the fact that several antimicrobial peptides are amyloid-like and *vice versa* amyloid peptides have antimicrobial properties. Both classes of peptides exhibit membrane-interaction and disruption ability with common mechanisms (*54, 55*). Interestingly, antimicrobial peptides and PLA_2_ enzymes can form amyloid-type co-fibrils that require the presence of the PLA_2_ lipid hydrolytic product, so the PLA_2_ catalytic activity synergizes with the propensity to form amyloid fibril (*56*). This can contribute to explain the synergy between catalytic and noncatalytic PLA_2_s, as observed for *B. asper* myotoxins I and II (*57*).

Why should secreted PLA_2_s form these complexes in membrane? Code et al (*27*) proposed that amyloid-type formation of PLA_2_ is functional to the control of enzyme action. We add that the function, on cell surface, could be also of triggering the endocytic process, only when protein concentration exceeds a threshold value. Phase transition in cell surface could be useful to select and concentrate secreted protein factors with consequent activation of a signaling cascade, membrane movements and internalization mechanisms. Different compositions of this multimolecular assembly could explain the various actions and specificities of secreted PLA_2_s.

In conclusion, with this work, we have identified, for the first time, a functional interaction between Mt-II and a cell surface protein, NCL. Our result definitively excludes that Mt-II interacts only with membrane lipids but also excludes interaction with a single protein receptor and indicates that this PLA_2_-like toxin, due to its propensity to form amyloid-like fibrils, probably participates in a multi-molecular assembly with many actors. We observed that secreted PLA_2_s possess prion-like sequences, in exposed loops, very similar to the low complexity domains interacting with NCL in phase transition phenomena inside the cell. We think therefore that the observed Mt-II/NCL protein assembly can be considered a phase separation on the cell surface, functional to the internalization of external factors. This internalization pathway, found for a snake toxin, may be the one followed by the human homologous protein, PLA2g2a, and this would explain its localization in nucleus and nucleolus. If so, it will be interesting to understand why so similar proteins, despite following the same path, cause such a different effect in cells.

## MATERIALS AND METHODS

### Isolation of Mt-II from *Bothrops asper* venom and modification with transglutaminase

Mt-II was isolated from the crude venom of *Bothrops asper,* a pool obtained from at least 30 specimens kept at the serpentarium of Instituto Clodomiro Picado, University of Costa Rica, as described in a previous work *(57).* The isolated Mt-II was modified by reaction with transglutaminase (TGase) from *S. mobaraensis* (Ajinomoto Co., Tokyo, Japan), as described in Spolaore et al. (2012)(?*8*), to obtain a protein mono-derivative with a fluorophore, for the imaging experiments, or with biotin for the pull-down experiments. Briefly, toxin was dissolved at a concentration of 1 mg/ml in 0.1 M sodium phosphate buffer (pH 7), to this solution the reactive Z-Gln-Gly-CAD-DNS, Z-Gln-Gly-CAD-TAMRA or Z-Gln-Gly-CAD-Biotine (ZEDIRA GmbH, Darmstadt, Germany) (stock solution 34 mg/mL in DMSO) was added at a protein/ligand molar ratio of 1/20. TGase was added at an enzyme/substrate (E/S) ratio of 1/25 (w/w), and the reaction mixtures were incubated for 4 hours at 25° C. Reactions were stopped by addition of iodoacetamide (100 μM final concentration). The fraction of mono-labeled Mt-II was purified from the reaction mixture by RP-HPLC with a C18 column (150 × 4.6 mm, 5 μm particle size; Phenomenex). The obtained product was lyophilized with a Freeze Dryer Edwards E2-MS (Milano) and analysed by a Q-Tof Micro mass spectrometer (Micromass, Manchester, UK). The toxic activity of the modified Mt-II was analyzed by a vitality test in RAW 264.7 cells and verified to be conserved (Fig. S1).

### Cell cultures, cytotoxicity and internalization assays

The murine macrophage cell line RAW264.7 (ATCC TIB 71) and skeletal muscle C2C12 (CRL-1772) were obtained from the American Type Culture Collection and were maintained in Dulbecco’s modified Eagle’s medium (DMEM) supplemented with heat-inactivated fetal bovine serum (EuroClone) 10%, 100 μg/ml streptomycin, 100 U/ml penicillin. Primary cultures of skeletal muscle cells were prepared from newborn (1-2 days-old) mice (CD1, Charles River) as reported by Massimino et al., 2006 *(58)* and seeded in Ham’s F12 (Eurobio) supplemented with 10% (v/v) fetal calf serum, 2 mM glutamine, 100 μg/ml penicillin, 100 μg/ml streptomycin (Eurobio). For C2C12 and primary myoblasts differentiation, the growth medium was replaced with DMEM containing 2% (v/v) horse serum (Eurobio) and the cells were incubated for 5-6 days. For the cytotoxicity assay macrophages and differentiated myotubes were grown in 96-well plates and then exposed to 2 μM Mt-II, with or without the indicated competitor, dissolved in modified Krebs-Ringer buffer (mKRB) (140 mM NaCl, 2.8 mM KCl, 2 mM MgCl_2_, 1 mM CaCl_2_, 10 mM Hepes, and 11 mM glucose, pH 7.4) for the indicated time. Cellular necrosis of C2C12 or primary mouse myotubes was estimated by measuring the release of LDH using the commercial kit TOX7 (Sigma). RAW264.7 cell death was evaluated with the colorimetric MTT assay, CellTiter96 (Promega). All antagonists were checked for their effect on cell viability and used at non-toxic concentrations. For the myotoxin internalization assay RAW264.7 cells and differentiated C2C12 or primary mouse myotubes grown in 96-well plates were incubated with 1 μM Mt-II-TAMRA in presence of different inhibitors for 10 minutes. The cells were then washed three times with ice-cold PBS to remove the non-internalized protein and lysed with 100 μL of lysis buffer (50 mm Tris, 0.8% Triton X-100, 0.2% SDS, pH 7.4). Cell-associated fluorescent protein was determined by measuring the fluorescence of the lysate using a TECAN Infinite M1000 microplate reader (Ex 547 nm and Em 573 nm). All data were collected in triplicate and three independent experiments were performed.

For Mt-II polymerization status analysis on primary myotubes (4•10^4^ cells cells/well p96, 6 differentiation days), cells were treated with 1 μM Mt-II for 30 min, 4°C. Then cells were washed with mKRB and treated (60 min, 4°C) with different quantities of the cross-linker BS3 (Thermo Scientific), resuspended in Laemmli Sample Buffer and analyzed by western blot (Bolt™ 4-12% Bis-Tris Plus Gels, Invitrogen) with conjugated streptavidin-HRP (1:1000, Invitrogen).

All aspects of animal care and experimentation were performed in compliance with European and Italian (D.L. 26/2014) laws concerning the care and use of laboratory animals. All experimental procedures and animal care protocols were approved by the Italian Ministry of Health, and by the Ethical Committee for animal care and use of the University of Padova (*OPBA, Organismo Preposto al Benessere degli Animali*).

### Pull-down experiments and cross-linking on magnetic beads

Biotinylated Mt-II (Mt-II-B, 2 μg) was combined with 5 μl of Pierce Streptavidin Magnetic Beads (Thermofisher), pre-washed according to the manufacturer’s instruction. Mt-II-B and beads were incubated for 1 hour in a 2 ml low binding tube (Eppendorf) in Thermomixer (Eppendorf) at 25°C, 500 rpm and then excess of Mt-II-B was washed with PBS. After each step of incubation or washing, beads were collected with a magnetic stand. Following steps of cell lysis were performed on ice, centrifugation was performed at 4 °C in a fixed angle rotor, incubations were performed in Thermomixer at 4°C, 500 rpm, equilibration and elution at 25°C, 500 rpm. RAW264.7 cells, seeded the day before in p6 (5 • 10^5^ /well), were washed with mKRB and lysed in 200 μl/well of breaking solution (BS) (50 mM TRIS, 25% (w/w) sucrose, 5% (v/v) glycerol, 5mM MgCl_2_, 2,8mM KCl, 2mM CaCl_2_ and Roche protease inhibitors) with 10 passages through a 25G needle. The obtained suspension was briefly vortexed and left on ice for 30 minutes and then centrifuged for 3 minutes to eliminate unbroken cells and most of the nuclei. The supernatant was collected and incubated overnight with 5 μl of streptavidin magnetic beads combined with Mt-II-B, or without Mt-II-B for background control. Then the beads were washed three times for 3 minutes each, in Thermomixer, with 200 jlll of BS and three times with 200 μl of IP Buffer (Pierce). The beads were finally equilibrated with three passages in 200 μl of 50 mM NH4HCO3 (pH 8) and the isolated proteins were eluted with 20 μl of 5% sodium deoxycholate (SDC) in 50 mM NH4HCO3. In the third experiment the elution step was preceded by crosslinking with 100 μl of 0.25 mM 3,3’-dithiobis(sulfosuccinimidyl propionate) (DTSSP, Thermo Scientific), in 50 mM NH_4_HCO_3_, at 25°C for 30 min. Reaction was stopped with 10 μl of 1 M TRIS pH 7.5 and beads were washed three times with 5% SDC in NH4HCO3 (at 25°C) to remove non-crosslinked proteins. Finally, elution was performed with 50 mM DTT in 5 % SDC, 50 mM NH4HCO3. Eluted proteins were analysed by electrophoresis in a Bolt™ 4-12% Bis-Tris Plus Gels (Invitrogen).

Membrane protein extracts were obtained with a Membrane Protein Extraction Kit (BioVision, Milpitas, CA) by following the manufacturer’s instructions, starting from 5 • 10^8^ RAW264.7 cells, 1.5 • 10^8^ C2C12 myotubes or 4.5 g of mouse posterior limb muscles (from 1 day-old mice). Membranes were resuspended in 1.2 ml of BS and pull-down was executed as reported in the previous paragraph, combining 200 μl of membrane suspension with 2 μg of Mt-II-B / 5 μl of streptavidin beads. Eluted proteins were analysed by western blot probed with anti-NCL rabbit polyclonal antibody C23 H-250 (Santa Cruz), diluted 1:200. BlueStar pLUS Prestained Marker (Nippon Genetics Europe GmbH) was used as MW standard. IRDYE 800cw, Goat anti-Rabbit (LI-COR)(1:10^4^) and IRDYE 680cw, Goat anti-Mouse were used as secondary antibodies. Pictures were acquired with Odyssey CLx (LI-COR) and analysed with Image Studio Software 4.0 (LI-COR). The kit efficiency was checked by western blot analysis of RAW264.7 subcellular fractions (see supplementary Fig. S2).

### Mass spectrometry analysis

Samples isolated from the pull-down experiments performed as described above were loaded in a 4-12% SDS-PAGE gel (NuPage, Invitrogen). The electrophoretic separation was stopped after about 10 min, as soon as the protein extracts completely entered the running gel. This preliminary step allows the removal of salts and any other possible interfering compounds from the sample. Gel bands were then excised, cut in smaller pieces, washed several times with 200 μL of 50 mM NH_4_HCO_3_ (pH=8) and dried under vacuum after a short wash with 200 μL of acetonitrile (ACN). Cysteines were reduced with 10 mM dithiothreitol (in 50 mM NH_4_HCO_3_) for 1 h at 56 °C, and alkylated with 55 mM iodoacetamide (in 50 mM NH_4_HCO_3_) for 45 minutes at room temperature (RT) in the dark. Gel pieces were then washed with alternate steps of 50 mM NH_4_HCO_3_ and ACN, and finally dried under vacuum. Proteins were in-gel digested with sequencing grade modified trypsin (Promega, Madison, WI, USA) at 37 °C overnight (12.5 ng/μL trypsin in 50 mM NH_4_HCO_3_). Peptides were extracted with three steps of 50% ACN/0.1% formic acid and the samples containing the peptide mixtures were dried under vacuum and stored at -20 °C until the LC-MS/MS analysis was performed.

LC-MS/MS analysis was conducted on an LTQ-Orbitrap XL mass spectrometer (ThermoFisher Scientific, Rockford, IL, USA), coupled with a nano-HPLC Ultimate 3000 (Dionex-ThermoFisher Scientific). Samples were loaded onto a pico-frit column (75 μm I.D., 15 μm tip, 11 cm length, New Objective) packed in house with C18 material (Aeris peptide 3.6 μm XB-C18, Phenomenex) and separated using a 45-minutes linear gradient of ACN/0.1% formic acid (from 3%-40% ACN), at a flow rate of 250 nL/min. To avoid any possible carry-over effect on the chromatographic column, after each sample an identical analysis was performed by injecting a blank. The analysis was performed in a data-dependent mode: a full scan at 60,000 resolution on the Orbitrap was followed by MS/MS fragmentation scans on the four most intense ions acquired with collision-induced dissociation (CID) fragmentation in the linear trap. Raw data files were analyzed with the software MaxQuant (Cox, J. and Mann, M. *Nat Biotechnol,* 2008, 26, pp 1367-72) against the mouse section of the Uniprot database (version 20141201), concatenated with a small database of the most common contaminant proteins found in proteomics experiments. Enzyme specificity was set to trypsin with up to 2 missed cleavages. The mass tolerance window was set to 20 ppm for parent mass and to 0.5 Da for fragment ions. Carbamidomethylation of cysteine residues was set as fixed modification and methionine oxidation as variable modification. Results were filtered to keep into account only proteins identified with a false discovery rate (FDR) less or equal to 0.01 and with at least four independent unique peptides sequenced with high confidence by MS/MS. The intensity parameter calculated by MaxQuant was used to compare the abundance of the proteins isolated from the pull-down in the presence of Mt-II with those obtained from the control experiment.

### Live imaging, immunofluorescence and Congo Red staining of Mt-II intoxicated cells

RAW264.7 cells (1•10^5^ cells/13 mm, 5•10^5^/24 mm) or C2C12 (3.5•10^4^/13 mm, 1.5•10^5^/24 mm) or primary myotubes (3 • 10^5^/13 mm, 1.5 • 10^6^/24 mm) were seeded onto 13 mm or 24 mm coverslips coated with collagen 0.1% in HCl 0.01M (only for C2C12 and primary myotubes). For internalization experiments (Fig. 1) mouse primary myotubes were pre-treated for 20 minutes (37°C, 5% CO_2)_ with MitoTrackerTM Green and Hoechst 33342 diluted, 1:2000 and 1:1000 respectively, in culture medium. Cells were treated with a 1 μM solution of Mt-2-TAMRA in mKRB for 30 min, 37°C, 5% CO_2_, washed 3 times with mKRB and observed with an epifluorescence microscope. For time-lapse experiments (Video S1-S3) C2C12 myotubes or RAW264.7 were washed with mKRB, and treated with 1 μM Mt-II-DNS. Images were acquired sequentially every 2 minutes with a LEICA CTR6000 epifluorescence microscope (λ_exc_= 405 nm, and transmitted light filtered with DIC Nomarski). Videos are composed by two frames for second.

For colocalization experiments with surface NCL, live RAW264.7 cells and myotubes were incubated with anti-NCL rabbit polyclonal antibody C23 H-250 (Santa Cruz) diluted 1/50 in Optimem (Gibco) for 45 minutes at RT, then treated with 1 μM Mt-II-TAMRA in Optimem for 15 min, at 4°C. After extensive washings in PBS, cells were fixed with PFA (2% in PBS) for 20 minutes 4°C and incubated for 1 h, RT with the secondary antibody Alexa Fluor 488-conjugated anti-rabbit IgG (1:500, Molecular Probes). For amyloid-like structures, after 1 μM Mt-II treatment, 10 min, 4°C, cells were stained for 20 minutes at RT with Congo Red (Sigma) 0.02 mg/ml in PBS and 0.01% NaOH. Cell nuclei were counter-stained with Hoechst 33342 (5 μg/ml, Sigma). Finally coverslips were mounted in 8% Mowiol 40-88 (Sigma) in glycerol and PBS (1:3 (v/v)].

Images were acquired with inverted fluorescence microscope (Leica CTR6000) equipped with a computer-assisted charge-coupled camera (Orca Flash 4.0, Hamamatsu), or with a confocal microscopy system (Leica TCS-SP5) and with DMR microscope (Leica) to evidence amyloid-like structures under cross-polarized light.

## ACKNOWLEDGMENTS

We thank Cesare Montecucco, Paola Caccin, and Silvio Tosatto for helpful discussion. Stefania Ferro for technical contribution in F3 peptide synthesis. Kim Capuzzo for experiments performed during her graduation thesis. Cassa di Risparmio di Padova e Rovigo (Cariparo) Holding for funding the acquisition of the LTQ-Orbitrap XL mass spectrometer. This work was supported by the Interomics project of the CNR (CM), the International Center for Genetic Engineering and Biotechnology (ICGEB; CRP/13/006), and Vicerrectoría de Investigación (UCR 741-B4-100 and UCR 741-B5-602)

### Author contributions

F.T., M.L.M. designed research; F.T., M.L.M. and M.S. performed research; B.S. performed and analysed transglutaminase reaction; C.F. and G.A. performed mass spectrometry analysis; O.M. F3 peptide synthesis. L.M., J.F., B.L. snake venom toxins purification and characterization; F.T. wrote the paper with the contribute of M.L.M., G.A., L.M., J.F., and B.L.

The authors declare no conflict of interest.

